# The multiscale topological organization of the functional brain network in adolescent PTSD

**DOI:** 10.1101/2023.12.21.572743

**Authors:** David Corredor, Shailendra Segobin, Thomas Hinault, Francis Eustache, Jacques Dayan, Bérengère Guillery-Girard, Mikaël Naveau

## Abstract

The experience of an extremely aversive event can produce enduring deleterious behavioral and neural consequences, among which posttraumatic stress disorder (PTSD) is a representative example. In this work, we aim to study the whole-cortex functional organization of adolescents with PTSD without the *a priori* selection of specific regions of interest or functional networks. To do so, we built on the network neuroscience framework and specifically on multisubject community analysis to study the functional connectivity of the brain. We show, across different topological scales (the number of communities composing the cortex), an increased coupling between regions belonging to unimodal (sensory) regions and a reduced coupling between transmodal (association) regions in the adolescent PTSD group. These results open up an intriguing possibility concerning an altered large-scale cortical organization in adolescent PTSD.

**Significance Statement:** The understanding of brain responses following the experience of a traumatic event and the eventual apparition of posttraumatic stress disorder (PTSD) symptoms during youth remains an active topic of research in clinical neuroscience. We adapted a multisubject community detection algorithm to study the functional brain organization of PTSD from a whole cortex and multiscale topological perspective, thereby taking into account the complex (network) organization of the brain. PTSD patients and control subjects presented a different whole cortex functional organization that remained present across topological scales. PTSD patients showed a decreased interaction in transmodal cortices and, inversely, an enhanced interaction in unimodal cortices. This investigation highlights the importance of studying the brain from a complex and whole cortex perspective.

## 1. Introduction

Non-invasive brain imaging techniques that allow the investigation of brain architecture and dynamics in vivo and in real-time have become a central and promising tool to understand the neural consequences of exposure to a potentially traumatic event. An open topic in clinical neuroscience is how, after experiencing an extreme aversive event, some people might develop (or exacerbate) a maladaptive cerebral and behavioral profile and the manifestation of posttraumatic stress disorder (PTSD) symptoms. Although definitive brain signatures are still missing, recent technological developments in brain imaging and sophistication in data-analysis pipelines have provided important results on the brain alteration accompanying trauma exposure and PTSD.

Initial results from animal models and early activation paradigms using symptom provocation or trauma-related images showed that the amygdala, the medial prefrontal cortex, and the hippocampus are critical brain regions in PTSD appearance and symptomatology (Pitman et al., 2012; Shin et al., 2006). Subsequently, with the increased use of the resting state paradigm and the discovery of the cerebral cortex organization in intrinsic networks (IN) (for review, see Uddin et al., 2022), the focus on the activation of specific regions was completed by large-scale brain analysis in PTSD (Ross & Cisler, 2020). Investigations of the connectivity profile between a region of interest (ROI) and the rest of the brain (i.e., seed-based analysis) showed an altered large-scale brain organization in PTSD involving regions from the default mode (DMN), salience (SN), and control networks (CN) (Ross and Cisler, 2020).

Seed-based analysis requires the *a priori* selection of a ROI and overlooks brain regions and connections outside the direct connections with it. However, it has been shown that the brain is a complex system (Avena-Koenigsberger et al., 2018; de Schotten & Forkel, 2022), meaning that its functional properties arise from the interaction among its constituents, including indirect connections between regions. In this vein, recent studies using a whole-cortex approach, without the *a priori* selection of ROIs, presented complementary results to the activation and seed-based approaches on PTSD (Breukelaar et al., 2021; Misaki et al., 2018; Shaw et al., 2022).

Importantly, exposure to extreme aversive events can arrive early in life, with a particular propensity during adolescence (McLaughlin et al., 2013). The high rate of trauma exposure during childhood and adolescence leading to the potential development of PTSD and the associated impairments in daily life requires the investigation of the neural consequences of experiencing a traumatic event during youth (Cisler & Herringa, 2021; Marshall, 2016). Similar to adult PTSD, the majority of investigations on adolescent PTSD have been conducted using activation or seed-based analysis showing alterations principally concerning brain regions in the DMN, SN, and CN and their putative cognitive reciprocals: episodic memory (stimuli-independent mental activity), salience detection, and executive functions, respectively (for a review, see Cisler and Herringa, 2021; Ross and Cisler, 2020).

Although in recent years, increased consideration has been paid to the neural specificities of trauma exposure and PTSD during adolescence (Leibenluft & Barch, 2021), to our knowledge, no study has investigated the whole-cortex organization in adolescent PTSD. This work aims to fill this gap. To do so, we built on the network neuroscience framework, specifically in community analysis (Betzel, 2020). We used the multisubject community detection (MSCD) method (Betzel et al., 2019) to compare the community organization in functional brain data between a group of non-exposed adolescents and a group of exposed adolescents presenting severe PTSD symptoms.

The MSCD approach allowed us to investigate the functional organization in PTSD at both the global, whole-cortex organization (i.e., the difference in the community organization) and the local pairwise connections between brain regions. Furthermore, by realizing a data-driven clustering of functional communities, we circumvent the overreliance on functional adult templates (e.g., the Yeo atlas; Yeo et al., 2011) to study the adolescent brain functional networks and move beyond the focus on the DMN, SN, and CN and the putative behavioral and cognitive categories they are related to.

## 2. Methods

We present a novel analysis and results using a clinical data set that has been previously published. Details of the participants, clinical assessments, and neuroimaging data acquisitions can be found in (Viard et al., 2019) and are briefly summarized below.

### 2.1 Experimental dataset and data acquisition

Fifteen adolescents with PTSD, aged 13 to 18 years old, were recruited. Data from one patient were unusable because she slept during the resting state fMRI scan. All presented chronic PTSD for at least six months. Twenty-four typically developing adolescents with no history of trauma were chosen to match the patient group regarding age and IQ. Altogether, 14 PTSD patients (12 females) and 24 controls (13 females) were included in the analyses (for more details, see Viard et al., 2019). The study was approved by the local Ethics Committee (CPP Nord Ouest III). All adolescents and their parents signed informed consent after a comprehensive description of the study.

All participants were scanned with a Philips Achieva 3.0 T MRI scanner at the Cyceron Center (Caen, France). High-resolution T1-weighted anatomical volumes were acquired using a three-dimensional fast-field echo sequence (3D-T1-FFE sagittal; repetition time=20ms, echo time=4.6ms, flip angle=10°, 180 slices, slice thickness=1mm, field of view=256×256mm^2^, in-plane resolution=1×1mm^2^). Resting-state fMRI functional volumes were obtained using an interleaved 2DT2* SENSE EPI sequence designed to reduce geometric distortions with parallel imaging (2D-T2∗-FFE-EPI axial, SENSE=2; repetition time=2382ms, echo time=30ms, flip angle=80°, 42 slices, slice thickness=2.8mm, field of view=224×224mm2, in-plane resolution=2.8×2.8mm2, 280 volumes). The resting state fMRI duration was 11.26 min.

### 2.2 Data pre-processing and subject functional co-activity network

Functional data were pre-processed using the CONN toolbox release 22.a (https://web.conn-toolbox.org/) and the Statistical Parametric Mapping toolbox (SPM 12, University College London, London, UK). Functional images were realigned to the first volume, slice-timing corrected, normalized into standard MNI space, and smoothed with a Gaussian filter kernel of 8 mm width half maximum (FWHM). Noise components from cerebral white matter and cerebrospinal areas, subject-motion parameters, scan outliers constant, and first-order linear session effects were entered as confounds in a first-level analysis. Acquisitions with framewise displacement above 0.9mm or global BOLD signal changes above 5 s.d were taken as noise components and used as potential confounding effects. Finally, a temporal band-pass filter (0.01–0.1 Hz) was applied. The pre-processed time series for each brain region was used to create the subject’s functional networks.

Here, we defined the network nodes using the 91 regions of the Harvard-Oxford parcellation, expanding the whole cortex. Furthermore, for their known relevance in PTSD pathology, we also included the bilateral amygdala and hippocampus from the sub-cortical Harvard-Oxford parcellation. We also ran the analyses using the Brainnetome parcellation to assess methodological robustness. The Brainnetome atlas comprises 246 regions, from which we selected all cortical regions plus the amygdala and hippocampus, resulting in 218 regions. Results and figures for the Brainnetome Atlas can be found in the supplementary document (Sup Figures 4 and 5).

Altogether, functional networks were summarized as adjacency matrices *W* = [*w*_*ij*_], *ij* ∈ [1,95] whose entries correspond to Z-transformed Pearson’s correlation coefficients between the pairwise time series of the 95 nodes. We kept signed coefficients without thresholding.

### 2.3 Community detection

Community detection applied to functional Magnetic Resonance Imaging (fMRI) is a data-driven method to identify brain communities based on the functional connectivity network of each subject (i.e., single-layer community detection; SLCD). In SLCD, a network is clustered by maximizing an objective modularity function by quantifying groups of nodes more connected to each other than expected in an appropriate null model (Betzel, 2020). To perform group comparisons, SLCD is applied to each subject’s functional connectivity matrix, allowing the subject information to be kept but at the risk of overfitting (Betzel et al., 2019). Otherwise, it is necessary to create an averaged connectivity matrix for each group and use SLCD to unravel its community structure, decreasing the risk of overfitting but blurring subject-specific information.

A method providing a trade-off between group-level and subject-level community detection comes from an extension of the SLCD dealing with multiple networks simultaneously (i.e., multilayer networks) (Mucha et al., 2010). Multilayer community detection has been applied extensively in neurosciences (Vaiana & Muldoon, 2020) and has recently been used in multilayer networks composed of functional connectivity matrices of different subjects (Betzel et al., 2019). In this way, a single structure (the multilayer network across subjects) contains all the information in the dataset while preserving subject-specific data.

### 2.4 Multisubject community detection

Similar to SLCD, multisubject community detection (MSCD) relies on the maximization of the modularity (Q) (equation 1). Therefore, as in the single-layer case, in MSCD, it is necessary to specify a null model and the resolution parameter (*γ*). Moreover, it is also required to precise the value of a second parameter controlling the interlayer strength coupling (*ω*) (eq.,1).

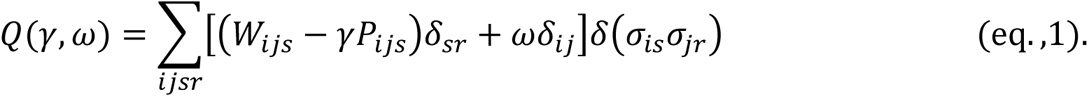

With:

– Q modularity quality metric
– *i,j* network nodes;
– *s,r* network layers;
– *W*_*ijs*_ weight of the edge linking nodes i and j in layer s;
– γ spatial resolution parameter (gamma);
– P_*ijs*_ weight of the edge linking nodes i and j in layer s in the null model;
– ω interlayer strength parameter (omega);
– δ_*s*r_ *=* 1 if *s* = r; 0 otherwise
– δ_*ij*_ *=* 1 if *i* = *j*; 0 otherwise
– σ_*is*_ community of node i in layer s
– δ(σ_*is*_σ_*j*r_) *=* 1 if σ_*is*_ = σ_*j*r_; 0 otherwise

Because we work with correlational networks, we use P_*ijs*_ = 1 as the null model in the modularity function (Bazzi et al., 2016; Betzel et al., 2015; Betzel et al., 2019).

For every iteration of the MSCD, we randomly selected 14 (among the 24) control subjects to build a balanced multilayer network regarding the number of subjects per group. Hence, the input of the MSCD is a multilayer network composed of 28 layers corresponding to the pairwise functional brain co-activity of 95 regions from the 14 PTSD subjects plus 14 control subjects (Figure 1, A).

**Figure 1.**
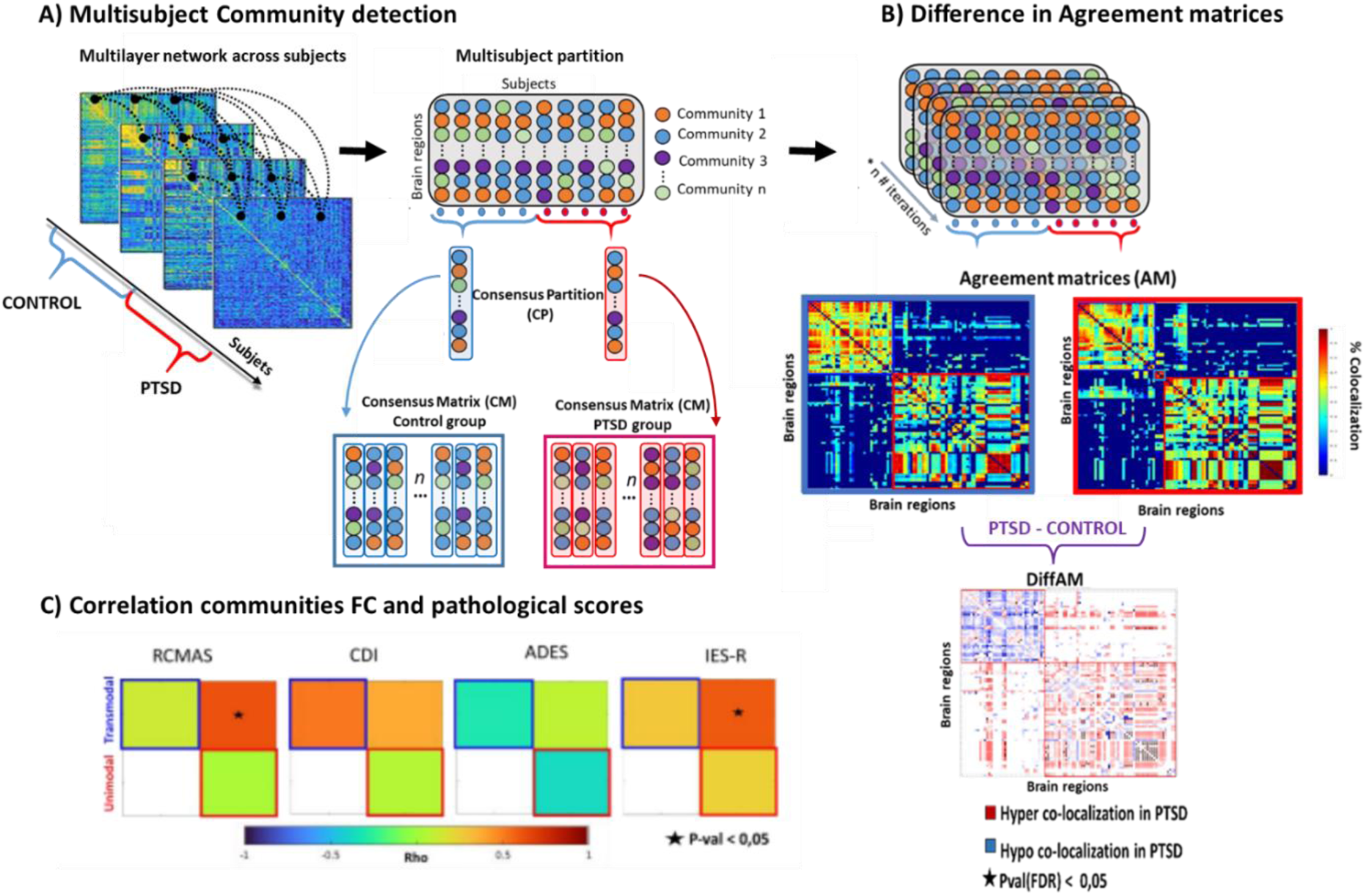
MSCD group comparison procedure. **A**) A multilayer network comprising 14 PTSD and 14 control subjects was subjected to a modularity maximization procedure to determine subjects’ partitions, summarized in the multisubject partition. For each group, we computed the consensus partition (CP) as the mode of a given region across subjects. To control for the degeneracy in the community detection algorithm and sweep across topological scales, we iterated this procedure multiple times. We computed the CPs for each iteration of the MSCD and computed the similarity of CPs across iterations for each group. **B)** Example for one topological scale: We calculated the Agreement Matrices (AM) representing the frequency at which any two nodes were assigned to the same community across subjects and iterations. We computed the difference between the AM of the two groups (DiffAM). In the DiffAM, entries > 0 (red) correspond to pairs of nodes that are more co-localized (hyper-colocalized) in the PTSD group compared to the control group. Entries < 0 (blue) correspond to hypo-colocalized nodes in the PTSD group compared to the control group, and entries equal to zero (green) denote no difference between groups. **C)** Finally, we went back to the functional connectivity (FC) matrices of PTSD subjects to calculate the correlation coefficients (rho) between FC and pathological scores (RCM AS: revised children’s manifest anxiety scale; CDI: Childhood depressive intervention; ADES: the adolescent dissociative experience; IES-R: the symptom severity scale).

### 2.5 Partition selection

After one iteration of the MSCD, we obtain a multi-subject partition matrix containing information about community membership for each node and subject in which rows correspond to brain regions and columns to subjects (Figure 1, A). The combination of *γ* and *ω* parameter values in the modularity function produces different partitions regarding the number of communities and the similarity of partitions across subjects. Thus, the landscape of possible multi-subject partitions can be organized according to the partitions’ topological scale (i.e., the mean *number of communities)* and *flexibility* (Betzel et al., 2019). In other words, multi-subject partitions can go from one single community to n communities (n = number of regions in the network) and have a flexibility value from 0 (partitions are identical across subjects) to 1 (partitions are totally different).

To run group comparisons, we start by filtering out multi-subject partitions obtained across iterations of the MSCD based on their flexibility and number of communities. First, we required comparable but not identical partitions across subjects to obtain a coherent group partition while keeping subject specificities. Thus, we excluded partitions with extreme flexibility values, keeping multisubject partitions with flexibility values within the range [0.2:0.8]. Second, based on Betzel et al. (2019), we excluded partitions with more than 20 communities, keeping partitions with a mean number of communities within the range [2:20], allowing us to investigate different topological scales of the brain functional organization.

We ran the MSCD algorithm with randomly selected pairs of *γ* and *ω* values until we obtained 10000 multisubject partitions within the previously defined flexibility and number of community boundaries (for details about these partitions, please see the supplementary information document and Sup Figure 1). Among this initially constrained set, we kept topological scales presenting the most stable organization (lower value in the average variation of information across partitions and iterations of the algorithm) for further analysis. Based on the variation of information (Sup Figure 2), we selected partitions composed of [2-4] communities (first topological scale), [5-7] communities (second topological scale), and [12-14] communities (third topological scale). Among the 10000 partitions, the first topological scale represents 1919, the second 3486, and the third topological scale 934 partitions.

**Figure 2.**
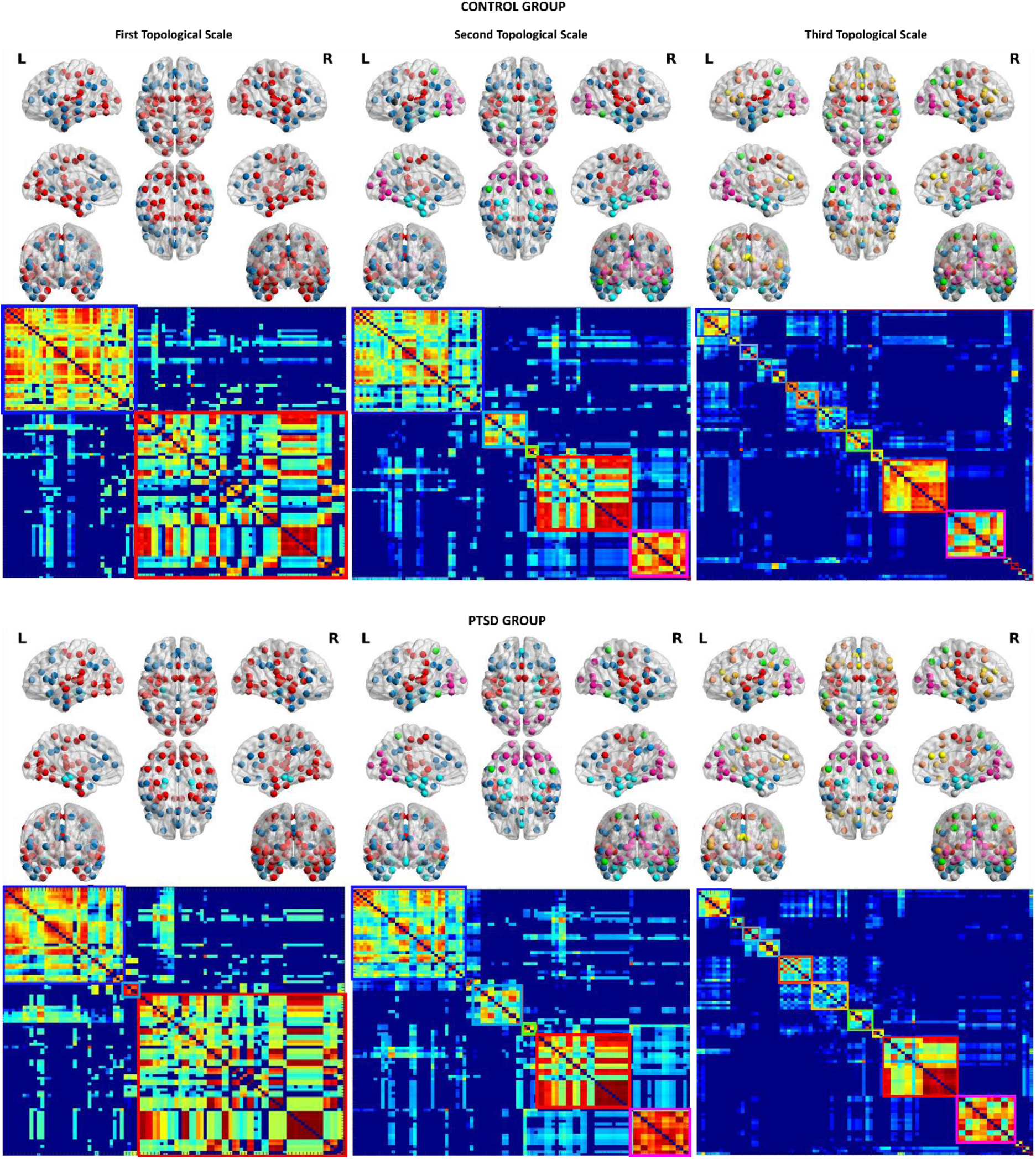
The whole-brain multiscale community organization. Representative partition of the control group (top) and PTSD group (bottom) for the first (left), second (middle), and third (right) topological scales. Colors in brain plots and boxes in agreement matrices denote community affiliation.

### 2.6. Communities’ analysis

For each iteration of the MSCD, a consensus partition is computed as the mode of community assignment across subjects for each brain region (Betzel et al., 2019) (Figure 1, A). By splitting the subjects according to group membership (i.e., control, PTSD), we obtained a consensus partition for each group. Then, for each topological scale, we summarized all iterations of the MSCD in a consensus matrix of dimensions [*n* × *i*], where n is the number of brain regions, and i is the number of partitions (Figure 1, A). We used the consensus matrix to test the representability of the partitions for each group. To do so, we assessed whether group partition similarities were higher within the control and PTSD groups than in two random groups constituted of a mixture of control and PTSD subjects (see section 2.8).

### 2.7. Pairwise analysis

To investigate the node-to-node specificities in brain organization, we compared the frequency at which any two nodes were allocated to the same community, summarized in the so-called Agreement matrix. In functional brain studies, it has been shown that agreement matrices contain critical information that is not accessible when solely studying functional connectivity matrices (Bassett et al., 2015).

For each group, we computed an agreement matrix (AM) summarizing information across all the partitions within the specific topological scale (Figure 1, B). The AM is a symmetric n*n matrix (n = 95 brain regions), representing the number of times, across the 14 subjects and the number of iterations, two nodes were assigned to the same community, divided by the maximum number of times they could be together. To control for spurious pairwise associations, we thresholded the AM matrices following the procedure in Bassett et al., 2013.

To compare the groups, we subtracted the AM of PTSD subjects from the AM of the control group (DiffAM) (Figure 1). In this new matrix, entries equal to zero denote no difference between groups. Entries > 0 correspond to pairs of nodes that are more co-localized (hyper-colocalized) in the PTSD group compared to the control group (Figure 1 red entries in DiffAM). Entries < 0 correspond to hypo-colocalized nodes in the PTSD group compared to the control group (Figure 1 blue entries in DiffAM).

### 2.8. Result significance

To compute statistics, we run 10000 times the analysis described in sections 1.6 and 1.7 using permuted versions of subjects’ group identities to create an empirical null distribution (Alexander-Bloch et al., 2012; Puxeddu et al., 2020). For each permutation, we reorganized the set of multisubject partitions according to a 1*28 vector of random subject labels (two groups of 14 subjects, each with a random mix of control and PTSD subjects). Results were corrected for multiple comparisons using false discovery rate (FDR) with a threshold of p<0.05.

### 2.9. Group partition and visualization

We computed the representative partition for each group and each topological scale corresponding to the consensus partition that presents the mean lower distance (higher similarity) among all consensus partitions (runs of the MSCD).

### 2.10. Correlation with pathological scores

We investigated whether the information in the unraveled communities was correlated with pathological scores associated with PTSD. To do so, we returned to the functional connectivity (FC) matrices of the PTSD participants and computed, for each topological scale, the within and between FC strength for each community in the representative partition of the PTSD group. We used screening tests of pediatric symptoms: the revised children’s manifest anxiety scale (RCMAS), Childhood depressive intervention (CDI), the adolescent dissociative experience (ADES), and the symptom severity scale (IES-R).

## 3. Results

### 3.1 Communities composition across topological scales

For both groups and atlases, the multiscale partitions were composed of distinct communities mirroring previous results concerning the fundamental brain’s functional organization. Figures 2 and 3 show the communities for the control and PTSD groups using the Harvard Oxford Atlas (partitions using the Braintome atlas can be found in the supplementary document, Figure 6). To better visualize the community organization in the third topological, see Figure 3.

**Figure 3.**
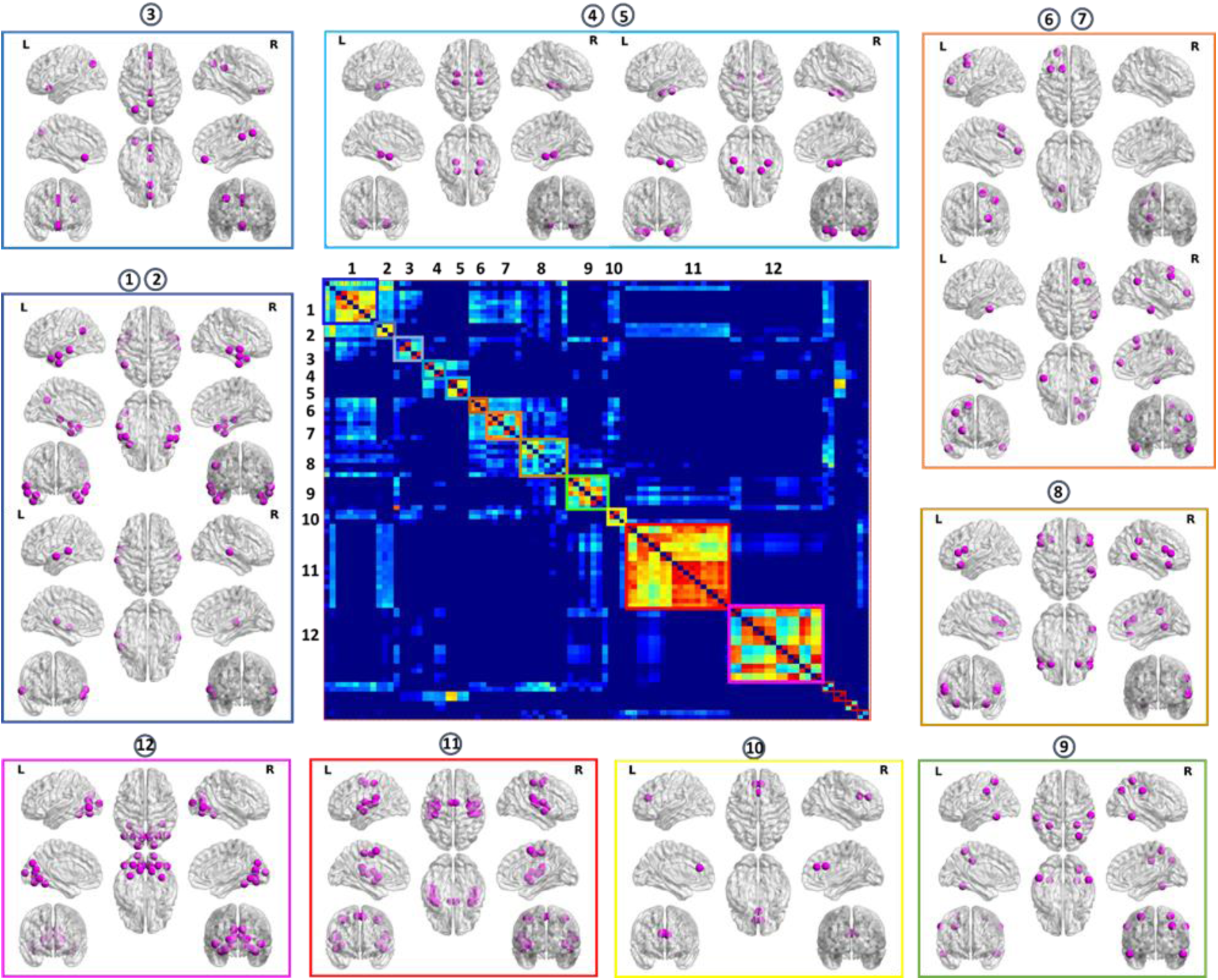
Agreement matrix and functional communities in the representative partition for the CONTROL group at the third topological scale (Harvard-Oxford Atlas; 95 regions). Box colors denote putative functional labels (see Table 1).

**Table 1.**
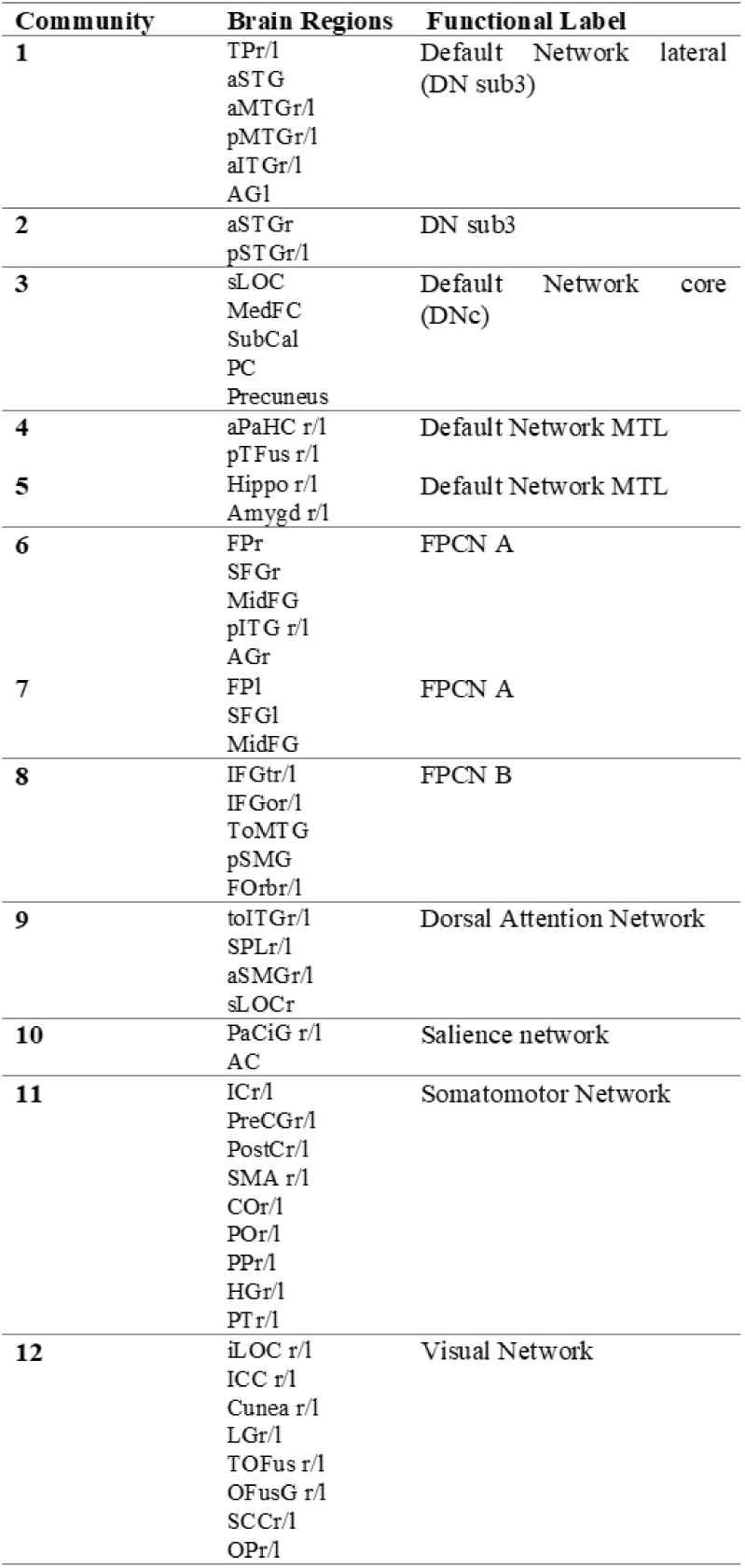
Labels of communities in the third topological scale for the control group. (see Figure 3). ‘r/l’ at the end of the region’s name denotes bilateral regions. R for right, l for left. Singletons’ communities’ (i.e., a one-node or bilateral node community) are not presented.

In the first topological scale (Figure 2, top left), the partition of the control group was composed of two communities. The first community (blue in Figure 2, top left) contains brain regions of the frontal, middle temporal, and posterior-anterior medial cortices. The second community (red in Figure 2, top) comprises occipital and pericentral brain regions. In the PTSD partition (Figure 2, bottom left), there was a third community comprising the hippocampal complex (light blue in Figure 2, bottom left).

For the second topological scale and the control group (Figure 2, top middle), the two communities in the first topological scale split into 5 communities corresponding to frontal and middle temporal cortices (blue in Figure 2, top middle), medial temporal regions (light blue), dorsal frontoparietal (green), pericentral (red), and occipital (pink), respectively. For the PTSD group, a small community comprising the precuneus and the posterior cingulate cortex appeared. Furthermore, the medial frontal and subcallosal cortex were allocated with the medial temporal regions and not in the frontal and lateral community, as in the control group.

Finally, in the third topological scale (Figure 2, top right), we found 12 communities (singletons excluded) containing medial frontoparietal (two communities, dark blue), medial anteroposterior (one community, blue), medial temporal (two communities, light blue), lateral frontoparietal (three communities orange), dorsal frontoparietal (green), midcingulo-insular (yellow), pericentral (red), the occipital (pink), and singletons communities (with single nodes; gray). For the PTSD group, we found 10 communities (singletons excluded) similar in composition to the control group.

### 3.2 Partitions are group dependent

We found a significant effect of group organization on the computation of group partitions for the first topological scale (p = 0.002), the second topological scale (p = 0.005), and the third topological scale (p = 0.001).

### 3.3. Hyper– and hypo-colocalized nodes in the PTSD group

To determine hyper-colocalized or hypo-colocalized nodes in the PTSD group compared to the control group, we used the difference between the two groups of the frequency at which any two nodes were allocated to the same community (Figure 1). We found a set of hyper-colocalized nodes (red edges in Figure 4 and supplementary Figures 7,8,9) in the occipital and pericentral communities for the three topological scales and the two atlases. Moreover, we found a set of hypo-colocalized nodes (blue edges in Figure 4 and Sup Figures 7,8,9) in the frontal, lateral, and medial anteroposterior communities.

**Figure 4.**
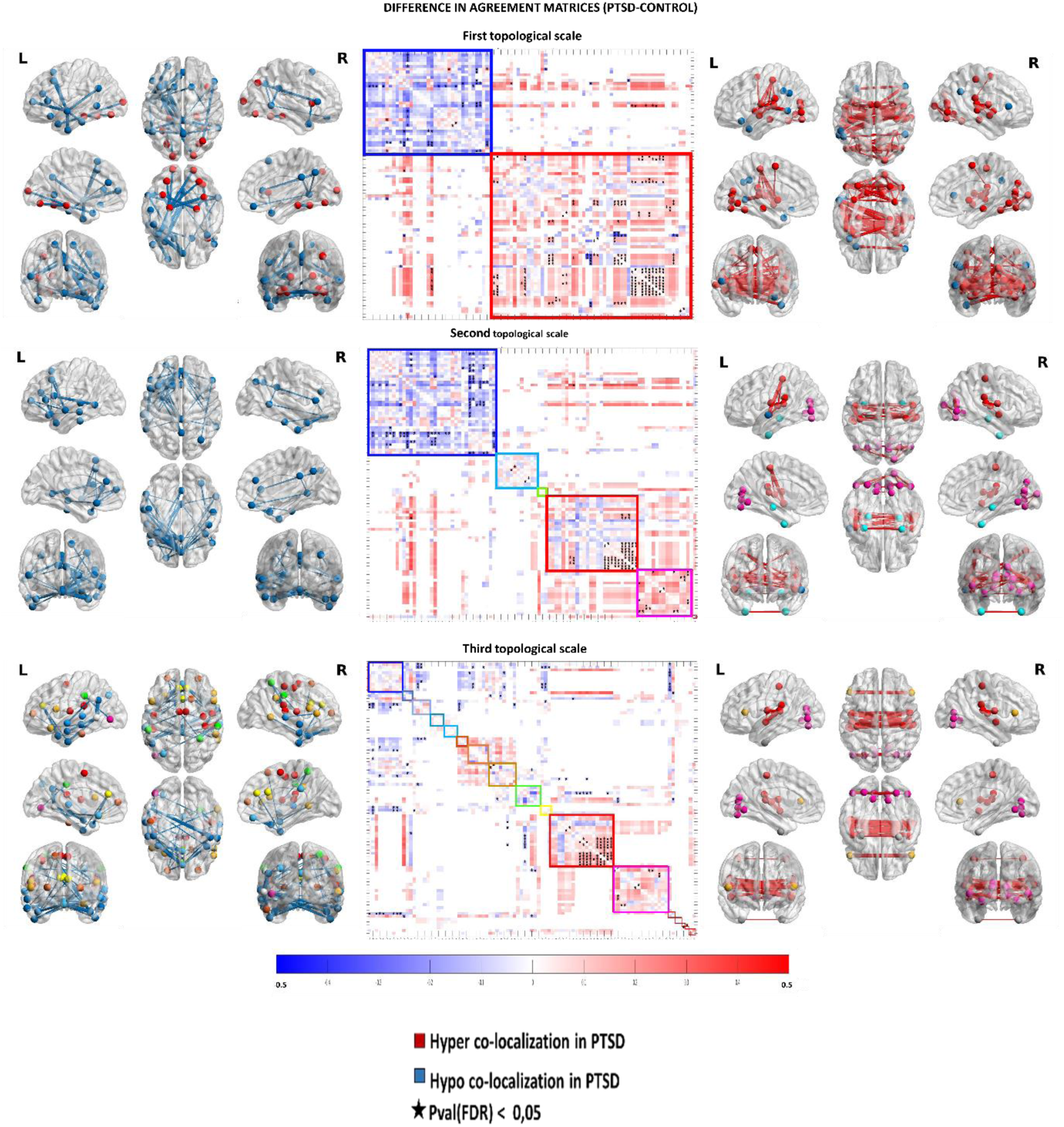
The difference in agreement matrices. (PTSD – Control) (DiffAM) in the first (top), second (intermediate), and third (bottom) topological scales (Harvard-Oxford Atlas). Entries > 0 (red) correspond to pairs of nodes that are more co-localized (hyper-colocalized) in the PTSD group compared to the control group. Entries < 0 (blue) correspond to hypo-colocalized nodes in the PTSD group compared to the control group, and entries equal to zero (white) denote no difference between groups. Entries with bright colors in the DiffAMs correspond to the remaining differences after permutations. Black stars in matrices and depicted edges in brain plots correspond to significant results after correction for multiple comparisons. For illustrative purposes, we plot the matrices according to the representative partition of the control group.

### 3.4 Correlation with pathological scores

After correction for multiple comparisons, no significant correlations were found between functional connectivity metrics and pathological scores. Please see the supplementary document to find the trends in the results.

### 3.5 Data and code availability

Subject-level adjacency matrices and code are available at https://doi.org/10.5281/zenodo.8164459.

## 4. Discussion

In the current study, we highlight population-specific characteristics of adolescent PTSD functional brain organization. Using the MSCD, we investigated the specificities in the network organization in adolescent PTSD across three topological scales at the global, whole-cortex organization, and local pairwise connections between brain regions.

### 4.1 An altered global organization in adolescent PTSD

For both control and PTSD groups, the multiscale partitions were composed of distinctive communities mirroring previous results concerning the fundamental brain’s functional organization in the unimodal-transmodal gradient (Huntenburg et al., 2018; Mesulam, 1998; Preti & Van De Ville, 2019; Shafiei et al., 2020; Vázquez-Rodríguez et al., 2019) on which specific brain regions (and intrinsic networks) are positioned (Margulies et al., 2016).

In the first topological scale, communities reflect a clear distinction between unimodal and transmodal regions (Figure 2, top). A transition from this bimodal organization towards the intrinsic network organization was observed at the second topological scale. Then, in the third topological scale, communities align with canonical intrinsic networks (INs) (Uddin et al., 2019) (Figure 3). See Table 1 and the supplementary document for a description of the nodes belonging to each community.

The fact that partitions were more similar within groups than between groups shows that partitions were representative of each group and supports the observation of a different global, whole-cortex, functional organization in adolescent PTSD compared to controls.

### 4.2 An altered pattern in the pairwise organization in adolescent PTSD

The differences observed when comparing the AM of the two groups (DiffAM) are spatially organized and remain present across topological scales. Furthermore, a more detailed picture emerges when carrying permutation analysis and controlling for multiple comparisons.

#### 4.2.1 Hypo-colocalized nodes in PTSD

Concerning the coarser topological scale (∼2 communities), we found a reduced coupling between regions in the “transmodal” (association) community in the PTSD group (Figure 4 top left, and Sup Fig 7). When increasing the number of communities (∼12 communities), the hypo-colocalized nodes were mainly within and between communities in association cortices containing nodes of the DMN (blue communities) and FPCN (orange communities). Importantly, we show that the decreased coupling of transmodal regions remains present across different topological resolutions. These results are in line with findings in adult and adolescent PTSD showing a decreased functional interaction between regions belonging to the DMN (Akiki et al., 2018; Bao et al., 2021; Miller et al., 2017; Sheynin et al., 2020; Viard et al., 2019). Our results also align with those of Breukelaar et al. (2021), where using a whole-brain approach in a cohort of adult PTSD, the authors found a hypo-connected subnetwork including regions from the DMN, control, and limbic (hippocampal complex + amygdala) communities in the PTSD group.

When running permutation analysis and controlling for multiple comparisons, we found the most implicated regions are in the anterior medial prefrontal cortex (amPFC) and the lateral-temporal cortex (black stars in Figure 4 left and sup Fig 4). The amPFC is a functional bridge between the DMNmtl and the DMNsub3, two subsystems composing the DMN (Andrews-Hanna et al., 2010; Christoff et al., 2016). Therefore, the hypo-colocalization found here could be related to a lack of global functional integration between DMN subsystems. Altogether, the decreased affiliation in DMN regions (maximally distant from sensory regions) (Margulies et al., 2016) and whose activity is mainly driven by endogenous dynamics and participating in the replay of past experiences (Kaefer et al., 2022) could be related to negative cognition and intrusive symptoms in PTSD patients.

#### 4.2.2 Hyper-colocalized nodes in PTSD

Across topological scales, we also found an increased coupling of regions in unimodal cortices (red and pink communities) (Figure 4, right). When controlling for spurious results using permutations and multiple comparisons, we found hyperconnected regions belonging to visual, auditory, and motor cortices.

In studies evaluating whole-brain resting-state functional connectivity in combat veterans, Misaki and colleagues (2018) found a decreased connectivity between the parahippocampal gyrus and occipital areas. In addition, Breukelaar et al. (2021) found regions from visual and somatomotor cortices belonging to a hypo-connected network. In the latter study, the participants were aged between 19 and 63 with a time since trauma of 0.75 to 51 years (about 20 years on average).

In the present study, the participants were younger with a more recent trauma, which could explain the still predominant involvement of the unimodal cortices. Váša and colleagues showed in healthy adolescents that, compared to transmodal regions, connectivity within sensory and motor cortices was stronger by the age of 14y. Importantly, there was an augmentation in sensory and motor connectivity strength across development from 14y to 26y (Váša et al., 2020). In our dataset, adolescents with PTSD were 15.7y old on average (1.51 sd), with an age of onset trauma of 13.3y (1.78 sd). Thus, our results might be related to the impact of experiencing a traumatic event in the initial condition of brain maturation in adolescence and their subsequent development. Future longitudinal studies are required to assess the impact of trauma on brain connectivity development during youth and early adulthood. Furthermore, unimodal cortices are implicated in the implementation of a defensive peripersonal space (defense zone with rapid detection of a potential threat) characterized by automatic protective actions (Rabellino et al., 2020). The enhanced connectivity in these regions could be thus related to hypervigilance-related symptoms in PTSD.

### 4.3 New avenues of research on PTSD

The results presented here open up an intriguing line of research to understand the underlying reasons for the exacerbated coupling in sensory regions and the decreased coupling in association regions. The unimodal-transmodal canonical organization in the functional macroscale brain architecture has been related to brain region characteristics, including i) the diversity in their time series, with sensory regions showing an increased pattern of dynamic diversity compared to association regions (Shafiei et al., 2020); ii) the coupling between the structural and functional connectivity, with sensory regions presenting an increased coupling strength between structure and function, compared to association cortices (Preti & Van De Ville, 2019; Vázquez-Rodríguez et al., 2019); iii) the temporal receptive windows (the time window within which a new stimulus will affect previously presented information), with sensory regions presenting a higher temporal integration (Chien & Honey, 2020; Hasson et al., 2015); and iv) the sensitivity of excitation-inhibition imbalance, with increased sensitivity in association regions (Yang et al., 2016).

Although this is not an exhaustive list accounting for the unimodal-transmodal organization, our results suggest that one or more of these characteristics could be at risk in patients with PTSD. Similar alterations to this organization have been shown in other clinical conditions, such as schizophrenia (Yang et al., 2016) and autism spectrum disorder (Watanabe et al., 2019). Thus, investigating the cortical gradient in PTSD appears as an interesting avenue to understanding the functional brain organization in this pathology and its relation to clinical symptoms.

Compared to other investigations studying the whole-brain functional organization in PTSD, our results present specificities that could be linked to adolescent PTSD compared to adult PTSD. Altogether, the increased coupling in sensorimotor regions could be a particularity of adolescent trauma, with an increased propensity to defensive strategies compared to adulthood. An alternative explanation accounting for these results would be linked to the proximity of the traumatic event rather than the age of trauma. Future longitudinal studies and direct comparisons between adult and adolescent PTSD are necessary to disentangle these two possibilities.

Importantly, this type of analysis could take advantage of an essential feature of the MSCD that we did not use here because of our small sample, namely the subject-specific community partitions. The possibility of obtaining reliable subject-specific partitions sharing community labels (e.g., community 1 in subject 1 is community 1 in subject 2) is a powerful tool that could be used in other investigations.

### 4.4 Methodological considerations

The main limitation of this work is the small sample size in our experiment. Therefore, the analysis presented here requires replication with a different and bigger cohort. Moreover, the absence of a control group of exposed subjects to a traumatic event without PTSD precludes a reliable conclusion concerning whether our results reflect PTSD rather than trauma-related specificities. Despite these issues, our investigation highlights the importance of paying attention to unexplored brain regions associated with sensorimotor processes and presents a method to compare whole-brain functional data from different populations straightforwardly.

From a methodological stance, MSCD relies on a specific definition of community, namely a set of nodes more connected to each other than with other nodes in the network (i.e., assortative communities). However, there are different possible modular organizations (e.g., core-periphery, disassortative) that are not studied when using the MSCD (Betzel, 2018; Fortunato & Hric, 2016; Hanteer & Magnani, 2020; Murphy et al., 2020)

## 5. Conclusion

In this work, we were interested in detailing the functional brain organization of adolescent PTSD using a whole cortex and multiscale topological approach. To do so, we built on the network neuroscience framework, specifically in multisubject community analysis. [] Our results open up an interesting perspective concerning a possible alteration in the large-scale cortical organization in PTSD. Testing for the possible explanations underlying the exacerbated coupling in sensory regions and the decreased coupling in association regions could be the object of future investigations.

## Authorship contribution

**David Corredor**: Conceptualization, Methodology, Formal analysis, Software, Writing – original draft. **Shailendra Segobin**: Conceptualization, Writing – review. **Thomas Hinault**: Writing – review. **Francis Eustache**: Funding acquisition, Writing – review. **Jacques Dayan**: Writing – review. **Bérengère Guillery-Girard**: Conceptualization, Supervision, Funding acquisition, Writing – review & editing. **Mikaël Naveau**: Conceptualization, Supervision, Writing – review & editing

## References

1. Akiki, T. J., Averill, C. L., Wrocklage, K. M., Scott, J. C., Averill, L. A., Schweinsburg, B., Alexander-Bloch, A., Martini, B., Southwick, S. M., Krystal, J. H., & Abdallah, C. G. (2018). Default mode network abnormalities in posttraumatic stress disorder: A novel network-restricted topology approach. NeuroImage, 176, 489–498. 10.1016/j.neuroimage.2018.05.005

2. Alexander-Bloch, A., Lambiotte, R., Roberts, B., Giedd, J., Gogtay, N., & Bullmore, E. (2012). The discovery of population differences in network community structure: New methods and applications to brain functional networks in schizophrenia. NeuroImage, 59(4), 3889–3900. 10.1016/j.neuroimage.2011.11.035

3. Andrews-Hanna, J. R., Reidler, J. S., Sepulcre, J., Poulin, R., & Buckner, R. L. (2010). Functional-Anatomic Fractionation of the Brain’s Default Network. Neuron, 65(4), 550–562. 10.1016/j.neuron.2010.02.005

4. Bao, W., Gao, Y., Cao, L., Li, H., Liu, J., Liang, K., Hu, X., Zhang, L., Hu, X., Gong, Q., & Huang, X. (2021). Alterations in large-scale functional networks in adult posttraumatic stress disorder: A systematic review and meta-analysis of resting-state functional connectivity studies. Neuroscience and Biobehavioral Reviews, 131, 1027–1036. 10.1016/j.neubiorev.2021.10.017

5. Bassett, D. S., Porter, M. A., Wymbs, N. F., Grafton, S. T., Carlson, J. M., & Mucha, P. J. (2013). Robust detection of dynamic community structure in networks. Chaos: An Interdisciplinary Journal of Nonlinear Science, 23(1), 013142. 10.1063/1.4790830

6. Bassett, D. S., Yang, M., Wymbs, N. F., & Grafton, S. T. (2015). Learning-induced autonomy of sensorimotor systems. Nature Neuroscience, 18(5), 744–751. 10.1038/nn.3993

7. Bazzi, M., Porter, M. A., Williams, S., McDonald, M., Fenn, D. J., & Howison, S. D. (2016). Community Detection in Temporal Multilayer Networks, with an Application to Correlation Networks. Multiscale Modeling & Simulation, 14(1), 1–41. 10.1137/15M1009615

8. Betzel, R. F. (2018). Diversity of meso-scale architecture in human and non-human connectomes. NATURE COMMUNICATIONS, 14.

9. Betzel, R. F. (2020). Community detection in network neuroscience. ArXiv:2011.06723 [q-Bio]. http://arxiv.org/abs/2011.06723

10. Betzel, R. F., Bertolero, M. A., Gordon, E. M., Gratton, C., Dosenbach, N. U. F., & Bassett, D. S. (2019). The community structure of functional brain networks exhibits scale-specific patterns of inter– and intra-subject variability. NeuroImage, 202, 115990. 10.1016/j.neuroimage.2019.07.003

11. Betzel, R., Mivsi’c, B., He, Y., Rumschlag, J., Zuo, X., & Sporns, O. (2015). Functional brain modules reconfigure at multiple scales across the human lifespan. ArXiv: Neurons and Cognition. https://www.semanticscholar.org/paper/Functional-brain-modules-reconfigure-at-multiple-Betzel-Mivsi’c/270044ccdace99af5e61b5f8cea72c8be253e36e

12. Breukelaar, I. A., Bryant, R. A., & Korgaonkar, M. S. (2021). The functional connectome in posttraumatic stress disorder. Neurobiology of Stress, 14, 100321. 10.1016/j.ynstr.2021.100321

13. Chien, H.-Y. S., & Honey, C. J. (2020). Constructing and Forgetting Temporal Context in the Human Cerebral Cortex. Neuron, 106(4), 675–686.e11. 10.1016/j.neuron.2020.02.013

14. Christoff, K., Irving, Z. C., Fox, K. C. R., Spreng, R. N., & Andrews-Hanna, J. R. (2016). Mind-wandering as spontaneous thought: a dynamic framework. Nature Reviews Neuroscience, 17(11), 718–731. 10.1038/nrn.2016.113

15. Cisler, J. M., & Herringa, R. J. (2021). Posttraumatic Stress Disorder and the Developing Adolescent Brain. Biological Psychiatry, 89(2), 144–151. 10.1016/j.biopsych.2020.06.001

16. Fortunato, S., & Hric, D. (2016). Community detection in networks: A user guide. Physics Reports, 659, 1–44. 10.1016/j.physrep.2016.09.002

17. Hanteer, O., & Magnani, M. (2020). Unspoken Assumptions in Multilayer Modularity maximization. Scientific Reports, 10(1), 11053. 10.1038/s41598-020-66956-0

18. Hasson, U., Chen, J., & Honey, C. J. (2015). Hierarchical process memory: memory as an integral component of information processing. Trends in Cognitive Sciences, 19(6), 304–313. 10.1016/j.tics.2015.04.006

19. Huntenburg, J. M., Bazin, P.-L., & Margulies, D. S. (2018). Large-Scale Gradients in Human Cortical Organization. Trends in Cognitive Sciences, 22(1), 21–31. 10.1016/j.tics.2017.11.002

20. Kaefer, K., Stella, F., McNaughton, B. L., & Battaglia, F. P. (2022). Replay, the default mode network and the cascaded memory systems model. Nature Reviews Neuroscience, 23(10), 628–640. 10.1038/s41583-022-00620-6

21. Lebois, L. A. M., Li, M., Baker, J. T., Wolff, J. D., Wang, D., Lambros, A. M., Grinspoon, E., Winternitz, S., Ren, J., Gönenç, A., Gruber, S. A., Ressler, K. J., Liu, H., & Kaufman, M. L. (2021). Large-Scale Functional Brain Network Architecture Changes Associated With Trauma-Related Dissociation. American Journal of Psychiatry, 178(2), 165–173. 10.1176/appi.ajp.2020.19060647

22. Leibenluft, E., & Barch, D. M. (2021). Adolescent Brain Development and Psychopathology: Introduction to the Special Issue. Biological Psychiatry, 89(2), 93–95. 10.1016/j.biopsych.2020.11.002

23. Margulies, D. S., Ghosh, S. S., Goulas, A., Falkiewicz, M., Huntenburg, J. M., Langs, G., Bezgin, G., Eickhoff, S. B., Castellanos, F. X., Petrides, M., Jefferies, E., & Smallwood, J. (2016). Situating the default-mode network along a principal gradient of macroscale cortical organization. Proceedings of the National Academy of Sciences, 113(44), 12574–12579. 10.1073/pnas.1608282113

24. Marshall, A. D. (2016). Developmental Timing of Trauma Exposure Relative to Puberty and the Nature of Psychopathology Among Adolescent Girls. Journal of the American Academy of Child and Adolescent Psychiatry, 55(1), 25–32.e1. 10.1016/j.jaac.2015.10.004

25. McLaughlin, K. A., Koenen, K. C., Hill, E. D., Petukhova, M., Sampson, N. A., Zaslavsky, A. M., & Kessler, R. C. (2013). Trauma exposure and posttraumatic stress disorder in a national sample of adolescents. Journal of the American Academy of Child and Adolescent Psychiatry, 52(8), 815–830.e14. 10.1016/j.jaac.2013.05.011

26. Mesulam, M. M. (1998). From sensation to cognition. Brain: A Journal of Neurology, 121 ( *Pt 6*), 1013–1052. 10.1093/brain/121.6.1013

27. Miller, D. R., Hayes, S. M., Hayes, J. P., Spielberg, J. M., Lafleche, G., & Verfaellie, M. (2017). Default Mode Network Subsystems Are Differentially Disrupted in Posttraumatic Stress Disorder. Biological Psychiatry: Cognitive Neuroscience and Neuroimaging, 2(4), 363–371. 10.1016/j.bpsc.2016.12.006

28. Misaki, M., Phillips, R., Zotev, V., Wong, C.-K., Wurfel, B. E., Krueger, F., Feldner, M., & Bodurka, J. (2018). Connectome-wide investigation of altered resting-state functional connectivity in war veterans with and without posttraumatic stress disorder. NeuroImage. Clinical, 17, 285–296. 10.1016/j.nicl.2017.10.032

29. Mucha, P. J., Richardson, T., Macon, K., Porter, M. A., & Onnela, J.-P. (2010). Community Structure in Time-Dependent, Multiscale, and Multiplex Networks. Science, 328(5980), 876–878. 10.1126/science.1184819

30. Murphy, A. C., Bertolero, M. A., Papadopoulos, L., Lydon-Staley, D. M., & Bassett, D. S. (2020). Multimodal network dynamics underpinning working memory. Nature Communications, 11(1). 10.1038/s41467-020-15541-0

31. Pitman, R. K., Rasmusson, A. M., Koenen, K. C., Shin, L. M., Orr, S. P., Gilbertson, M. W., Milad, M. R., & Liberzon, I. (2012). Biological studies of posttraumatic stress disorder. Nature Reviews Neuroscience, 13(11), 769–787. 10.1038/nrn3339

32. Puxeddu, M. G., Faskowitz, J., Betzel, R. F., Petti, M., Astolfi, L., & Sporns, O. (2020). The modular organization of brain cortical connectivity across the human lifespan. NeuroImage, 218, 116974. 10.1016/j.neuroimage.2020.116974

33. Rabellino, D., Frewen, P. A., McKinnon, M. C., & Lanius, R. A. (2020). Peripersonal Space and Bodily Self-Consciousness: Implications for Psychological Trauma-Related Disorders. Frontiers in Neuroscience, 14. https://www.frontiersin.org/articles/10.3389/fnins.2020.586605

34. Ross, M. C., & Cisler, J. M. (2020). Altered large-scale functional brain organization in posttraumatic stress disorder: A comprehensive review of univariate and network-level neurocircuitry models of PTSD. NeuroImage. Clinical, 27, 102319. 10.1016/j.nicl.2020.102319

35. Shafiei, G., Markello, R. D., Vos de Wael, R., Bernhardt, B. C., Fulcher, B. D., & Misic, B. (2020). Topographic gradients of intrinsic dynamics across neocortex. ELife, 9, e62116. 10.7554/eLife.62116

36. Shaw, S., Terpou, B., Densmore, M., Theberge, J., Frewen, P., McKinnon, M., & Lanius, R. (2022). Large-Scale Functional Hyperconnectivity Patterns Characterizing Trauma-Related Dissociation: A rs-fMRI Study of PTSD and its Dissociative Subtyp. 10.21203/rs.3.rs-2178523/v1

37. Sheynin, J., Duval, E. R., Lokshina, Y., Scott, J. C., Angstadt, M., Kessler, D., Zhang, L., Gur, R. E., Gur, R. C., & Liberzon, I. (2020). Altered resting-state functional connectivity in adolescents is associated with PTSD symptoms and trauma exposure. NeuroImage: Clinical, 26, 102215. 10.1016/j.nicl.2020.102215

38. Shin, L. M., Rauch, S. L., & Pitman, R. K. (2006). Amygdala, medial prefrontal cortex, and hippocampal function in PTSD. Annals of the New York Academy of Sciences, 1071, 67–79. 10.1196/annals.1364.007

39. Thomas Yeo, B. T., Krienen, F. M., Sepulcre, J., Sabuncu, M. R., Lashkari, D., Hollinshead, M., Roffman, J. L., Smoller, J. W., Zöllei, L., Polimeni, J. R., Fischl, B., Liu, H., & Buckner, R. L. (2011). The organization of the human cerebral cortex estimated by intrinsic functional connectivity. Journal of Neurophysiology, 106(3), 1125–1165. 10.1152/jn.00338.2011

40. Uddin, L., Betzel, R., Cohen, J., Damoiseaux, J., De Brigard, F., Eickhoff, S., Fornito, A., Gratton, C., Gordon, E., Laird, A., Larson-Prior, L., McIntosh, A., Nickerson, L., Pessoa, L., Pinho, A., Poldrack, R., Razi, A., Sadaghiani, S., Shine, J., & Spreng, R. N. (2022). Controversies and current progress on large-scale brain network nomenclature from OHBM WHATNET: Workgroup for HArmonized Taxonomy of NETworks. 10.31219/osf.io/25za6

41. Uddin, L. Q., Yeo, B. T. T., & Spreng, R. N. (2019). Towards a Universal Taxonomy of Macro-scale Functional Human Brain Networks. Brain Topography, 32(6), 926–942. 10.1007/s10548-019-00744-6

42. Vaiana, M., & Muldoon, S. F. (2020). Multilayer Brain Networks. Journal of Nonlinear Science, 30(5), 2147–2169. 10.1007/s00332-017-9436-8

43. Váša, F., Romero-Garcia, R., Kitzbichler, M. G., Seidlitz, J., Whitaker, K. J., Vaghi, M. M., Kundu, P., Patel, A. X., Fonagy, P., Dolan, R. J., Jones, P. B., Goodyer, I. M., the NSPN Consortium, Vértes, P. E., & Bullmore, E. T. (2020). Conservative and disruptive modes of adolescent change in human brain functional connectivity. Proceedings of the National Academy of Sciences, 117(6), 3248–3253. 10.1073/pnas.1906144117

44. Vázquez-Rodríguez, B., Suárez, L. E., Markello, R. D., Shafiei, G., Paquola, C., Hagmann, P., van den Heuvel, M. P., Bernhardt, B. C., Spreng, R. N., & Misic, B. (2019). Gradients of structure–function tethering across neocortex. Proceedings of the National Academy of Sciences, 116(42), 21219–21227. 10.1073/pnas.1903403116

45. Viard, A., Mutlu, J., Chanraud, S., Guenolé, F., Egler, P.-J., Gérardin, P., Baleyte, J.-M., Dayan, J., Eustache, F., & Guillery-Girard, B. (2019). Altered default mode network connectivity in adolescents with posttraumatic stress disorder. NeuroImage. Clinical, 22, 101731. 10.1016/j.nicl.2019.101731

46. Watanabe, T., Rees, G., & Masuda, N. (2019). Atypical intrinsic neural timescale in autism. ELife, 8, e42256. 10.7554/eLife.42256

47. Yang, G. J., Murray, J. D., Wang, X.-J., Glahn, D. C., Pearlson, G. D., Repovs, G., Krystal, J. H., & Anticevic, A. (2016). Functional hierarchy underlies preferential connectivity disturbances in schizophrenia. Proceedings of the National Academy of Sciences, 113(2), E219–E228. 10.1073/pnas.1508436113

