## Supplemental Information Appendix for "The multiscale topological organization of the functional brain network in adolescent PTSD"

#### SI Methods

_Partitions selection:_

An important concern when using modularity maximization to unravel brain communities is the selection of free parameters in the modularity function. In the case of multilayer networks, there are two free parameters to specify: The $\gamma$ parameter controls the number and size of communities, and the $\omega$ parameter dictates the interlayer connection's strength. Because there is no preferential number of communities or an ideal interlayer coupling value, selecting these parameters relies on heuristics, for instance, aiming to find the parameters providing the more stable partitions across iterations of the algorithm (Akiki & Abdallah, 2019).

In the present work, we aimed to select the modularity maximization parameters in a principled, data-driven way. We were interested in partitions that are comparable but not identical across subjects and allow interpretation based on previous studies in terms of INs. As in Betzel et al. 2019, we defined a large range of possible values for $\gamma$ and $\omega$. Then, a randomly selected pair of parameter values from the previously defined range was used to launch one run of the MSCD. We repeated this procedure until we obtained 10000 partitions of interest: partitions in which the difference in the community assignment of regions across subjects (their flexibility) and the number of communities were within reasonable but large boundaries. We retained multisubject partitions with a flexibility value within the range [0.2,0.8] (meaning that partitions are neither identical nor totally different across subjects) and a mean number of communities within the range [2,20].

Details of the $\gamma$, $\omega$, number of communities, and flexibility of the 10000 partitions can be found in supplementary figures 1 (Harvard-Oxford atlas) and 2 (Brainnetome atlas).

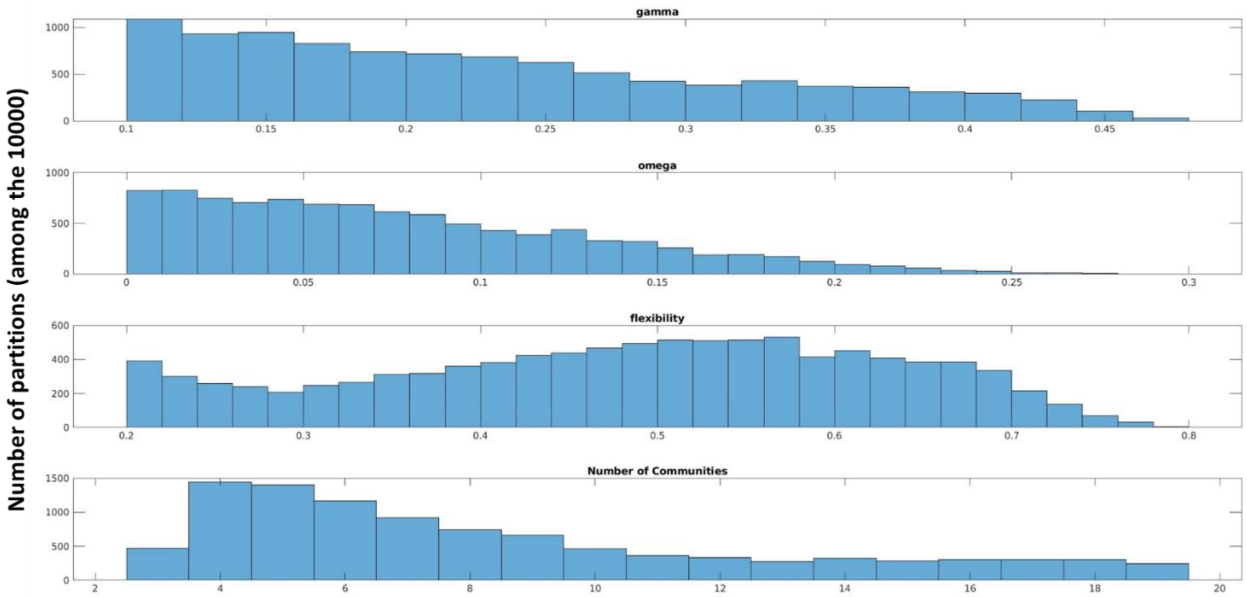


**Sup Figure 1. Gamma, omega, flexibility, and number of communities across the 10000 initial partitions for the Harvard-Oxford Atlas**

Among the initial 10000 partitions, we searched for the more stable topological scales (i.e., number of communities) based on a similarity metric. Because of its rapid computational implementation, we used the normalized variation of information (VIn). The rationale was to define the topological scales containing the more similar partitions across runs of the MSCD. To do so, for each group and topological scale, we computed the consensus partition's (dis)similarity values across runs of the MSCD (see *methods in* the main text) (supplementary Figures 2 and 5).

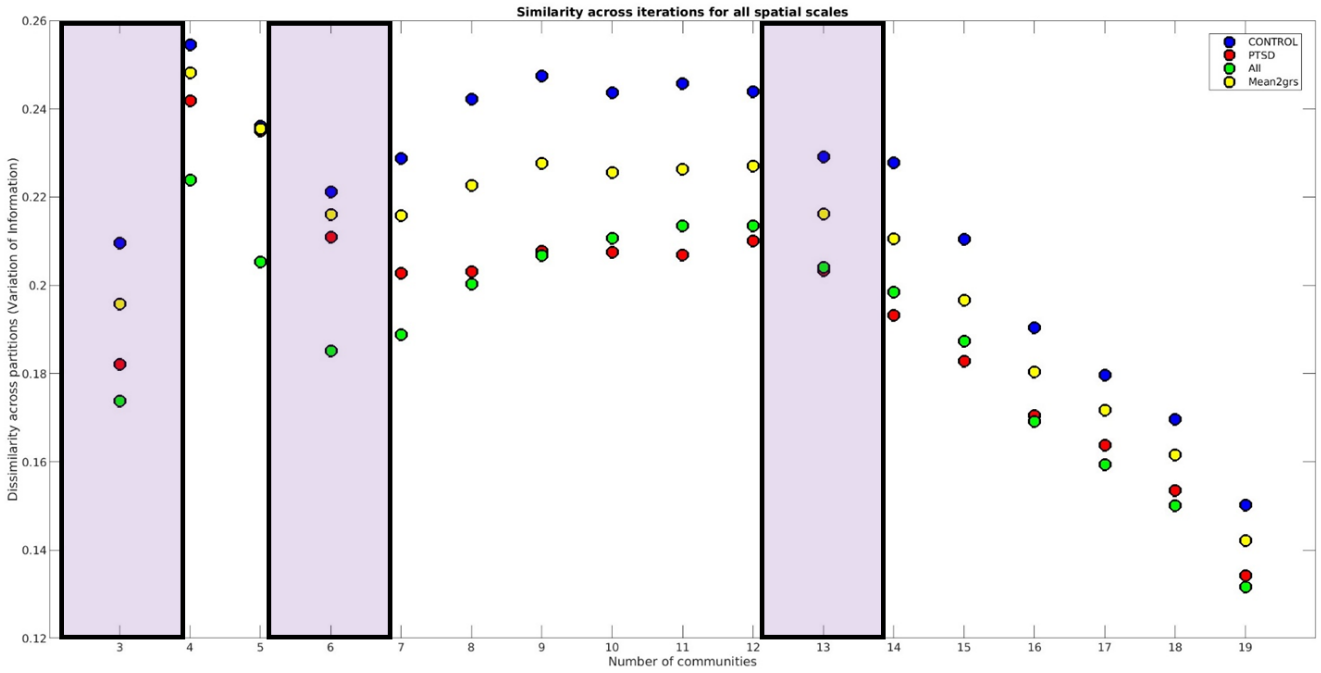


**Sup Figure 2. Consensus partition's (dis)similarity values across runs of the MSCD for the Harvard-Oxford Atlas. We selected for further analysis the set of partitions belonging to the spatial scales presenting the lower dissimilarity (higher similarity) across runs of the MSCD. Purple squares denote the selected topological scales.**

*Communities' composition and INs:*

In the main text (Figures 2 and 3), we give functional labels to the unraveled communities. The first topological scale partition comprised two communities dividing association to sensory regions. Concerning the third topological scale of the Harvard-Oxford atlas, we found 12 communities for the control group. Five communities were in line with the so-called Default Network (DN) (Andrews-Hanna et al., 2010; Christoff et al., 2016): Community 1 and 2 (dark blue) contained nodes from the medial temporal gyruses, inferior temporal gyruses, and the temporoparietal lobule (lateral DN; DNsub3). Community 3 (blue) included the medial frontal cortex, the subcallosal cortex, the posterior cingulate cortex, and the precuneus (anterior-posterior DN; DNcore). Finally, communities 4 and 5 (light blue) with nodes of the medial temporal lobe (DNmtl).

We found three communities in line with previous studies distinguishing the frontoparietal control network (FPCN) into two different subsystems (Dixon et al., 2018; Kam et al., 2019). Community 6 and 7 (dark orange) were composed of the middle frontal gyrus, the superior frontal gyrus, the frontal pole, and the posterior inferior temporal gyrus (FPCN$_A$); and community 8 (dark orange) with the inferior frontal gyrus, the posterior supramarginal gyrus and the medial temporal gyrus (FPCN$_B$).

Community 9 (green) comprises the superior parietal lobule nodes, anterior supramarginal gyrus, and the inferior temporal gyrus aligning with the Dorsal Attention Network, DAN. Community 10 (yellow) comprises the anterior cingulate and bilateral paracingulate cortex (Salience). Community 11 (red), with precentral and postcentral gyri and the auditory cortex regions (Somatomotor-auditory). Community 12 (pink) with occipital nodes (Visual). Finally, three communities were composed of singletons nodes.

*Correlation with pathological scores:*

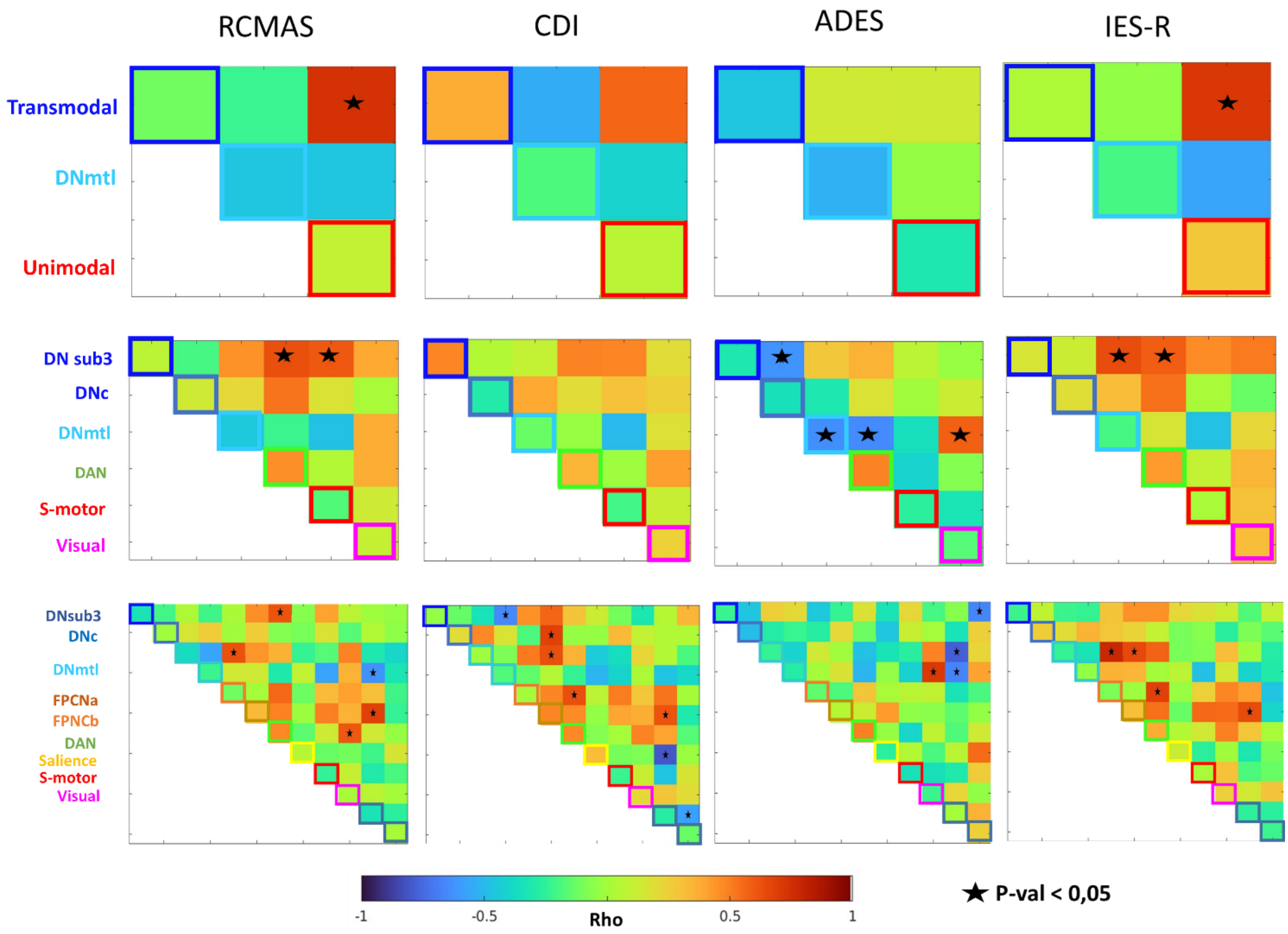


**Sup figure 3. Correlation between functional connectivity and pathological scores for PTSD subjects.** Top, first topological scale. Intermediate, second topological scale, and bottom, third topological scale. RCMAS: the revised children's manifest anxiety scale, CDI: Childhood depressive intervention, ADES: the adolescent dissociative experience, IES-R: the symptom severity scale.

### ***Analysis replication using the Brainntome atlas***

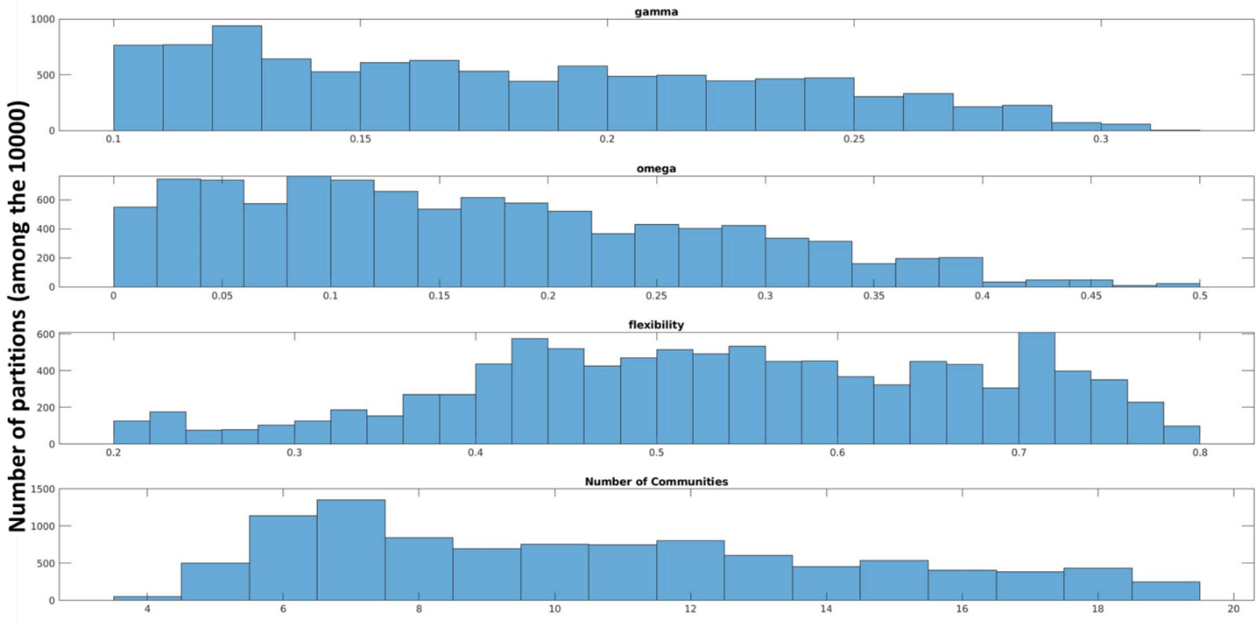


**Sup Figure 4. Gamma, omega, flexibility, and number of communities across the 10000 initial partitions for the Brainntome Atlas**

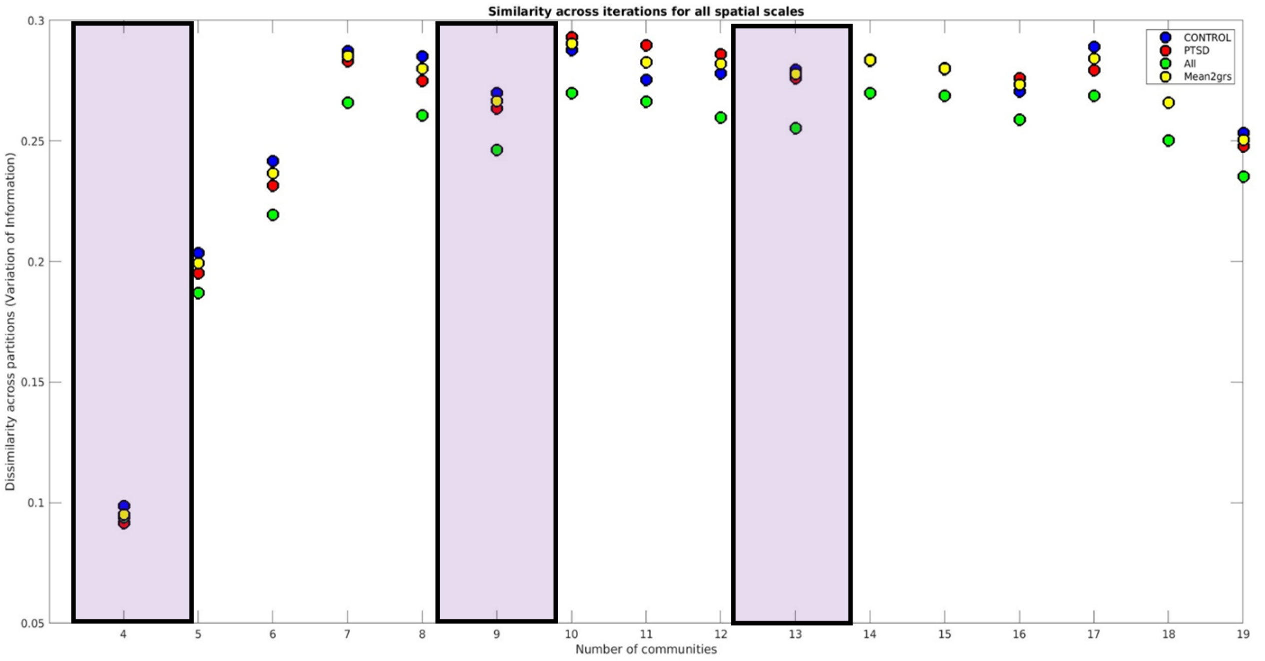


**Sup Figure 5. Consensus partition's (dis)similarity values across runs of the MSCD for the Brainntome Atlas. We selected for further analysis the set of partitions belonging to the spatial scales presenting the lower dissimilarity (higher similarity) across runs of the MSCD. Purple squares denote the selected topological scales.**

**Partition selection Brainntome Atlas**

For the Brainntome Atlas, we select partitions composed of [3-5] communities (first topological scale), [8-10] communities (second topological scale), and [12-14] communities (third topological scale). The first topological scale comprised 1698 partitions, the second 2300 partitions, and the third topological scale with 1874 partitions.

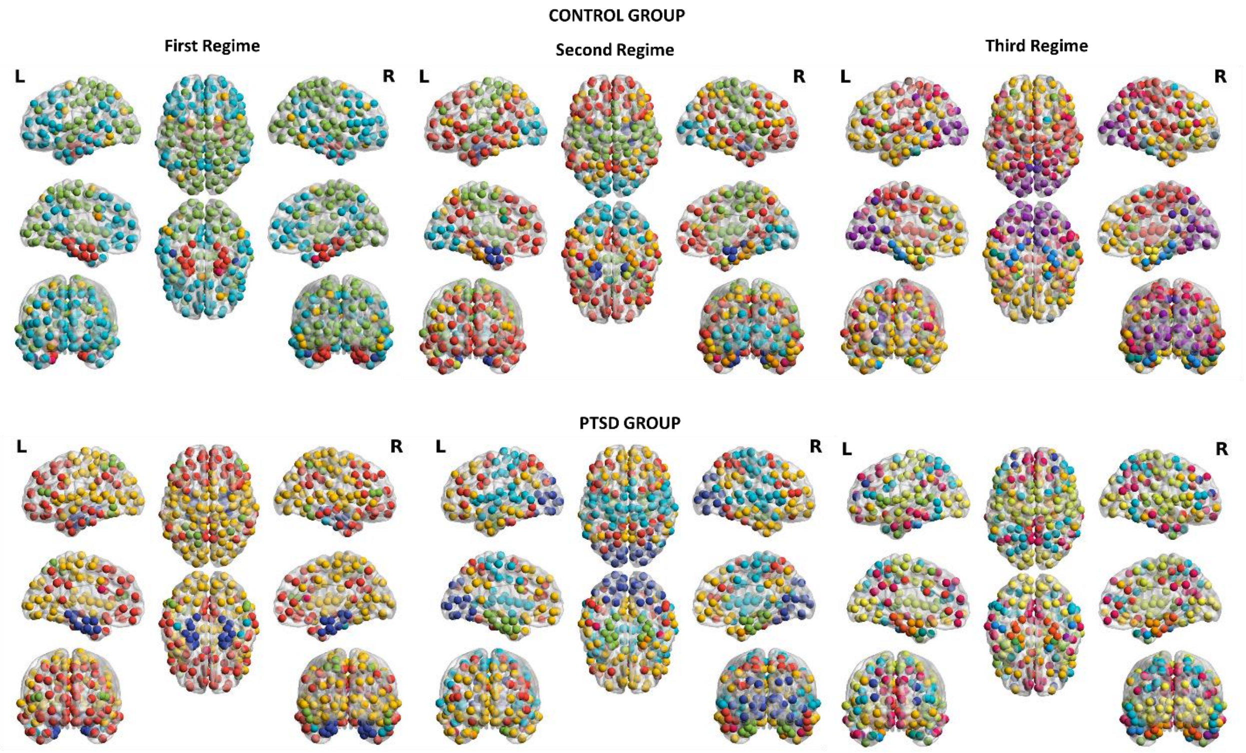


**Sup Figure 6. Brain plots of the functional communities in the representative partition for the control (top) and the PTSD (bottom) groups (Brainntome Atlas; 218 regions). Colors denote nodes' community membership.**

*<u>Replication for the Brainntome atlas: Partitions are group dependent for</u>*

We found a significant effect of group organization on the computation of group partitions for the first topological scale (p = 0.02), the second topological scale (p = 0.03), and the third topological scale (p = 0.01).

*Hyper- and hypo-colocalized nodes in the PTSD group:*

We found the same pattern of results when using the Brainntome Atlas. A set of hypo-colocalized nodes in association cortices, particularly in the DN, and a set of hyper-colocalized nodes in sensory cortices.

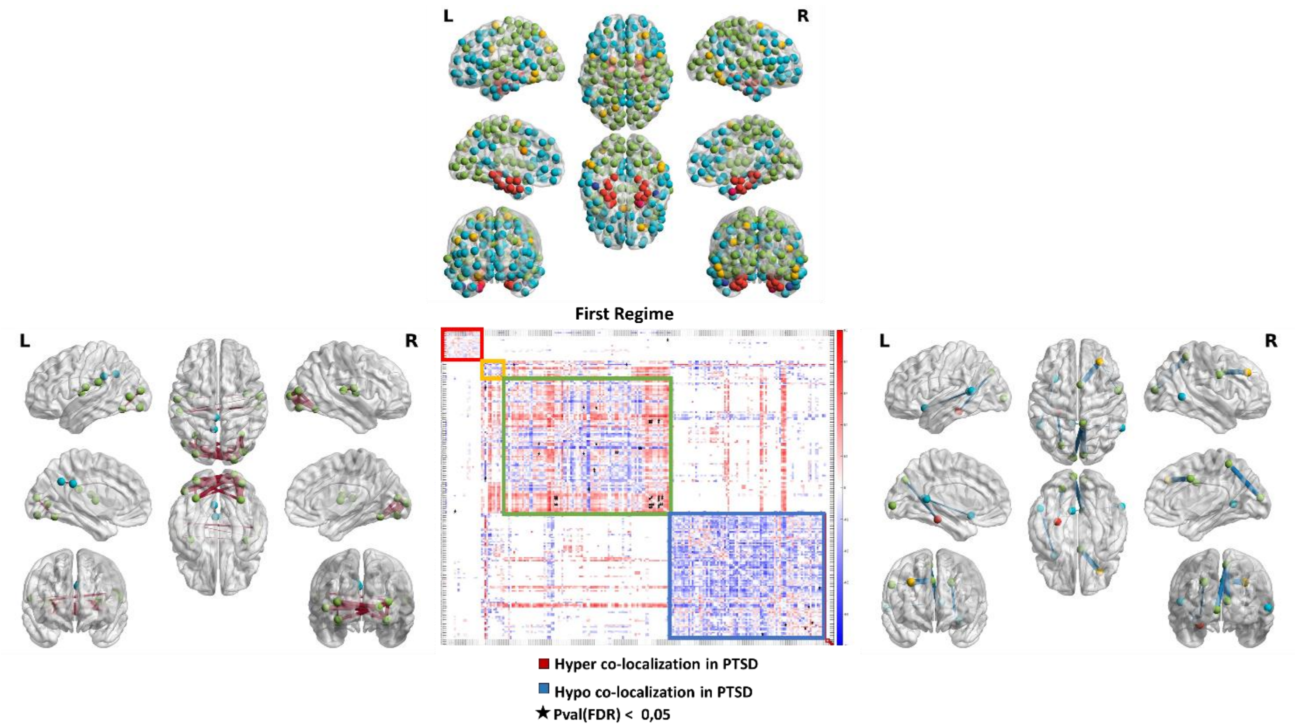


**Sup Figure 7. The difference in agreement matrices (PTSD – Control) in the first topological scale (Brainntome Atlas). Set of significant hyper-colocalized (red) or hypo-colocalized (blue) nodes in the PTSD group. Box colors in the matrix correspond to community colors in the brain plots. For illustration purposes, we show the partition of the representative control group.**

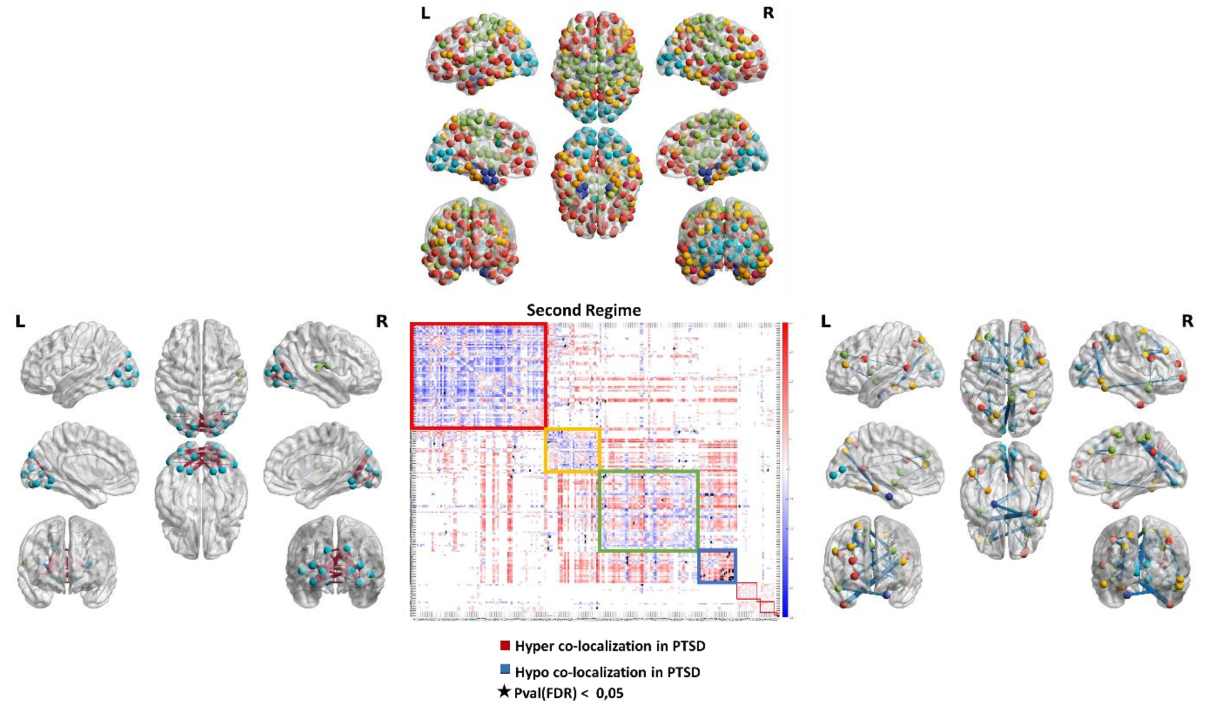


**Sup Figure 8. The difference in agreement matrices (PTSD – Control) in the second topological scale (Brainntome Atlas). Set of significant hyper-colocalized (red) or hypo-colocalized (blue) nodes in the PTSD group. Box colors in the matrix correspond to community colors in the brain plots. For illustration purposes, we show the partition of the representative control group.**

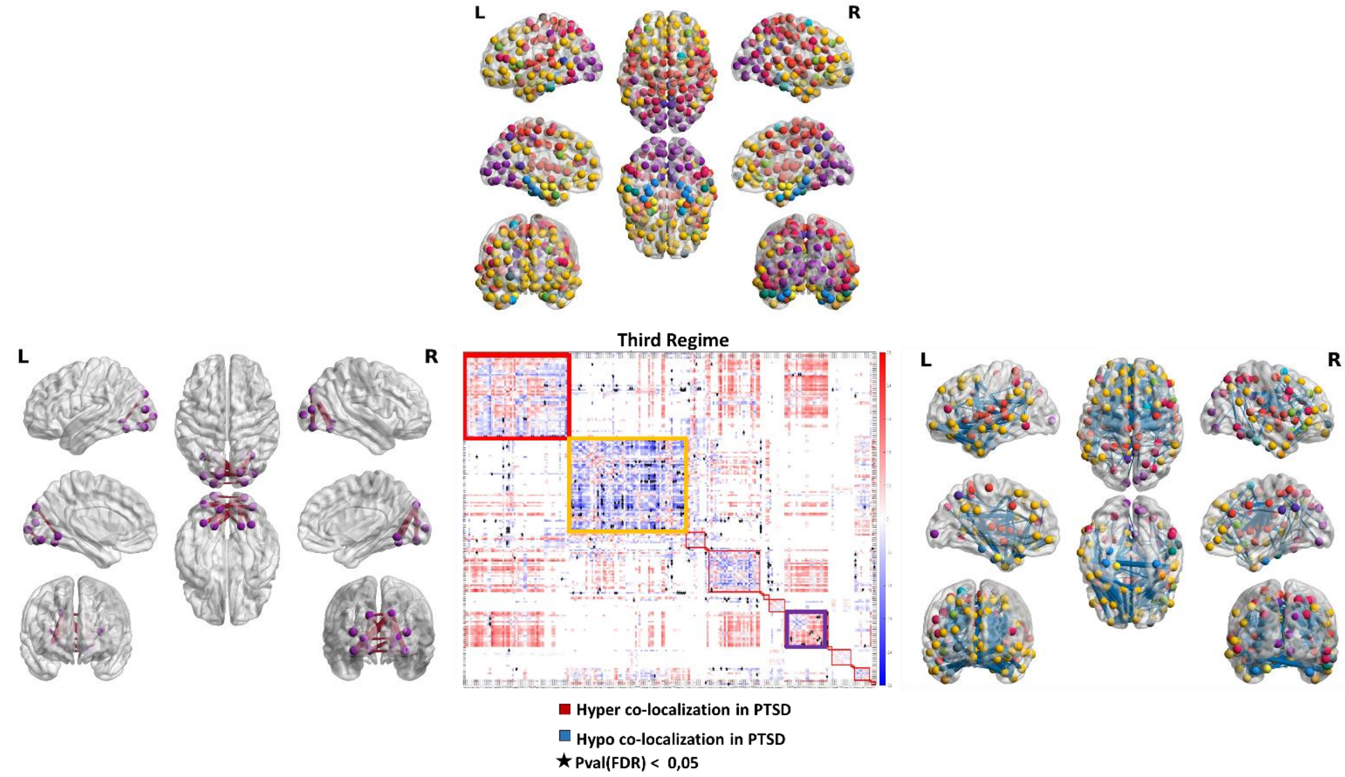


**Sup Figure 9. The difference in agreement matrices (PTSD – Control) in the third topological scale (Brainntome Atlas). Set of significant hyper-colocalized (red) or hypo-colocalized (blue) nodes in the PTSD group. Box colors in the matrix correspond to community colors in the brain plots. For illustration purposes, we show the partition of the representative control group.**

**REFERENCES**


Akiki, T. J., & Abdallah, C. G. (2019). Determining the Hierarchical Architecture of the Human Brain Using Subject-Level Clustering of Functional Networks. *Scientific Reports*, *9*(1). https://doi.org/10.1038/s41598-019-55738-y

Andrews-Hanna, J. R., Reidler, J. S., Sepulcre, J., Poulin, R., & Buckner, R. L. (2010). Functional-Anatomic Fractionation of the Brain's Default Network. *Neuron*, *65*(4), 550-562. https://doi.org/10.1016/j.neuron.2010.02.005

Christoff, K., Irving, Z. C., Fox, K. C. R., Spreng, R. N., & Andrews-Hanna, J. R. (2016). Mind-wandering as spontaneous thought: a dynamic framework. *Nature Reviews Neuroscience*, *17*(11), 718-731. https://doi.org/10.1038/nrn.2016.113

Dixon, M. L., De La Vega, A., Mills, C., Andrews-Hanna, J., Spreng, R. N., Cole, M. W., & Christoff, K. (2018). Heterogeneity within the frontoparietal control network and its relationship to the default and dorsal attention networks. *Proceedings of the National Academy of Sciences*, *115*(7). https://doi.org/10.1073/pnas.1715766115

Kam, J. W. Y., Lin, J. J., Solbakk, A.-K., Endestad, T., Larsson, P. G., & Knight, R. T. (2019). Default network and frontoparietal control network theta connectivity supports internal attention. *Nature Human Behaviour*, *3*(12), 1263-1270. https://doi.org/10.1038/s41562-019-0717-0
